# Detection of *Anopheles pseudopunctipennis* (Diptera: Culicidae) in Cochabamba, Bolivia, after 90 years of suspected presence

**DOI:** 10.64898/2025.12.05.692495

**Authors:** Libia Torrez, Lineth Garcia, Silvia Castellón, Rubén Castillo, Frédéric Lardeux

## Abstract

**Objectives.:** This study provides the first confirmed record of the malaria vector *Anopheles pseudopunctipennis* in Cochabamba city, Bolivia, ending nearly 90 years of suspected presence since the malaria epidemics of the 1930s and 1940s. Additionally, it assesses the potential suitability of the region for this mosquito species.

**Methods:** Mosquito larvae were collected as part of a comprehensive survey of Culicidae in Cochabamba city and its suburbs. Larvae of *An. pseudopunctipennis* were collected on April 16, 2024, and reared to the fourth instar in the laboratory. One male specimen obtained from collected larvae was pinned. Specimens were morphologically identified using a standard identification key and species re-description. The larval identification was based on its unique spiracular apparatus, while adults were identified by their characteristic black hind tarsomeres and unspotted wing costa. The specimens are preserved in the Laboratory of Medical Entomology at UMSS, Cochabamba.

**Results:** A single larval habitat was identified in Lincoln Park, in the center of Cochabamba. It consisted of a 184 m² pond with a cement bottom, clear water with very slow flow from a small spring, and abundant *Rhizoclonium* algae. A Maxent habitat suitability model indicated that the Cochabamba Valley provides suitable environmental conditions for *An. pseudopunctipennis*.

**Conclusions:** This study provides the first concrete evidence of *An. pseudopunctipennis* in Cochabamba city, confirming its presence after decades of speculation. However, its low abundance and limited larval habitats due to urbanization and pollution suggest that it poses no significant malaria transmission risk in the area.

## Introduction

*Anopheles pseudopunctipennis* is a mosquito species recognized as a significant vector of *Plasmodium vivax*, a malaria parasite affecting humans, in the mountainous regions of Mexico, Central America, and the Andean countries of South America, including Bolivia. In Bolivia, this species is found across a wide altitudinal range, from 300 to nearly 3000 meters ^1^. The Bolivian National Vector Control Program has been controlling the mosquito population through indoor insecticide spraying campaigns since the 1960s ^2^ to combat malaria transmission. Currently, it is acknowledged that malaria has almost been eradicated from the Andes region, with no malaria epidemics recorded since a small outbreak in 1998 ^3^, and except for some isolated cases that may occasionally appear ^4^. In the Cochabamba Department, autochthonous cases now appear to be rare and absent in the urban area of Cochabamba city ^5^.

However, malaria epidemics were recorded in the Cochabamba department during the 1930s and 1940s . The Cochabamba Valley was recognized as a hotspot for malaria transmission, with the city of Cochabamba facing a large epidemic in 1940-41.

Neighboring localities, such as Parotani and Quillacollo, also experienced outbreaks in 1936 ^2^. No vector species was identified at that time, but it is reasonable to suspect *An. pseudopunctipennis*, as it is the only *Anopheles* species present in the Andes area of Bolivia above 1000 meters of altitude, besides *An. argyritarsis*, which is not a vector ^6^.

Since the beginning of malaria concerns in the urban area of Cochabamba and neighboring localities, there have been no records of any *Anopheles* species, despite historically documented epidemics. More recently, surveys conducted in Cochabamba city by the Escuela Técnica de Salud de Cochabamba (Rodriguez, pers. comm.) and the Medical Entomology Laboratory of the Universidad Mayor de San Simón (LEMUMSS) identified seven mosquito species, all belonging to the genera *Culex* and *Aedes,* but no *Anopheles* ^7^.

This article presents the first documented record of *An. pseudopunctipennis* in Lincoln Park, located in the center of Cochabamba city, Bolivia, and provides insights into the potential suitability of the region for the species through Maxent modeling.

## Study area

The discovery of *An. pseudopunctipennis* occurred during surveys conducted as part of the research project DB.07 of the Medical Entomology Laboratory of the Universidad Mayor de San Simón, titled: *’Estudio de la fauna de culícidos vectores de enfermedades en el eje metropolitano de Cochabamba*’ ^8^. The project, which follows on from earlier laboratory studies ^7^, encompassed the Municipalities of Cochabamba Cercado, Colcapirhua, Quillacollo, Sacaba, and Tiquipaya, and was aimed at updating the list of the Culicidae fauna of the area, and identifying potential disease vectors. These municipalities are situated in the Central Valley Basin of the department, which is a semi-arid valley at an average altitude of 2,550 meters above sea level. The climate is temperate, with an average annual temperature of 20°C. The rainy season lasts approximately six months, from mid-October to mid-April, with a peak period of 31 days of rainfall exceeding 13 mm, centered around mid-January. The dry season lasts about six months, from mid-April to mid-October. The entire basin has a population of approximately 1.2 million inhabitants, distributed between highly urbanized areas, which account for 60% of the surface area, and agricultural zones. The region also has one of the highest population growth rates in the country. The area is experiencing rapid urbanization, with urban development encroaching on rural territories and leading to a significant loss of rural zones.

The potential larval habitats for mosquito larvae in the Cochabamba region are diverse, primarily due to the availability of various water resources. These habitats include rivers and their edges, lagoons, wetlands, irrigation channels, and during the rainy season, roadside ditches and small depressions that can turn into breeding grounds. The region’s rivers, including the Rocha River, Mailancu River, Huayculi River, La Tamborada River, and several northeastern streams, share a similar hydrological regime characterized by sudden, short-duration floods during the rainy season and drastically reduced flow during the dry season. Due to the high level of urbanization in the area, water sources also include domestic and industrial discharges. Despite their contamination, these waters are used for crop irrigation, leading to significant environmental issues. Natural, industrial and domestic water resources are all channeled for irrigation purposes to various agricultural areas such as Molinos, Chilimarka, Putuku, Linde, Canarancho, Villa Esperanza, Chiquicollo, Barrio Flores, Coña Coña, Sirpita, Rumi Mayu, Cuatro Esquinas, and Kallajchullpa. Additionally, there are domestic and peridomestic larval habitats, such as water storage containers, discarded tires, and other artificial containers found in residential areas. These water sources and irrigation channels, along with domestic and peridomestic habitats, create ideal conditions for the development of mosquito larvae.

## Methods

### Larval mosquito sampling

The DB.07 project basically consisted in the sampling of mosquito larval habitats for collecting the Culicidae fauna and registering basic ecological information. In the field, mosquito larvae were collected using the standard entomological dipping technique ^9^. Essential ecological data for each collected point, including GPS coordinates, types of larval habitats, and water quality were collected and organized using a cellular phone equipped with the VECTOBOL database ^10^. The database was designed with REDCap electronic data capture tools ^11, 12^ hosted at IRD-France. Then, in the laboratory, once species identifications were completed for each sample, the database was updated.

### Morphological identification of *An. pseudopunctipennis*

The collected larvae were brought to the laboratory and reared to the L4 stage. They were then subjected to a standard slide mounting process, which involved clearing in 10% KOH, an acetic acid bath, successive alcohol baths with increasing concentrations from 70% to 100%, a bath in eugenol, and mounting in Euparal. One male was also obtained from the single larva collected on March 26, 2024 and was pinned.

*Anopheles pseudopunctipennis* was morphologically identified and differentiated from other *Anopheles* using a standard identification key ^13^ and the species re-description ^14^. The larva is readily identified by “its spiracular apparatus that has two posterolateral spiracular lobe plates each with elongate, slender, sclerotized projection or "tail" from inner caudal margin” ^14^ (Figure 1). It is the only *Anopheles* species with such a characteristic. Adults are also easily identified by their hind tarsomeres entirely black and the wing costa without white spots.

**Figure 1.**
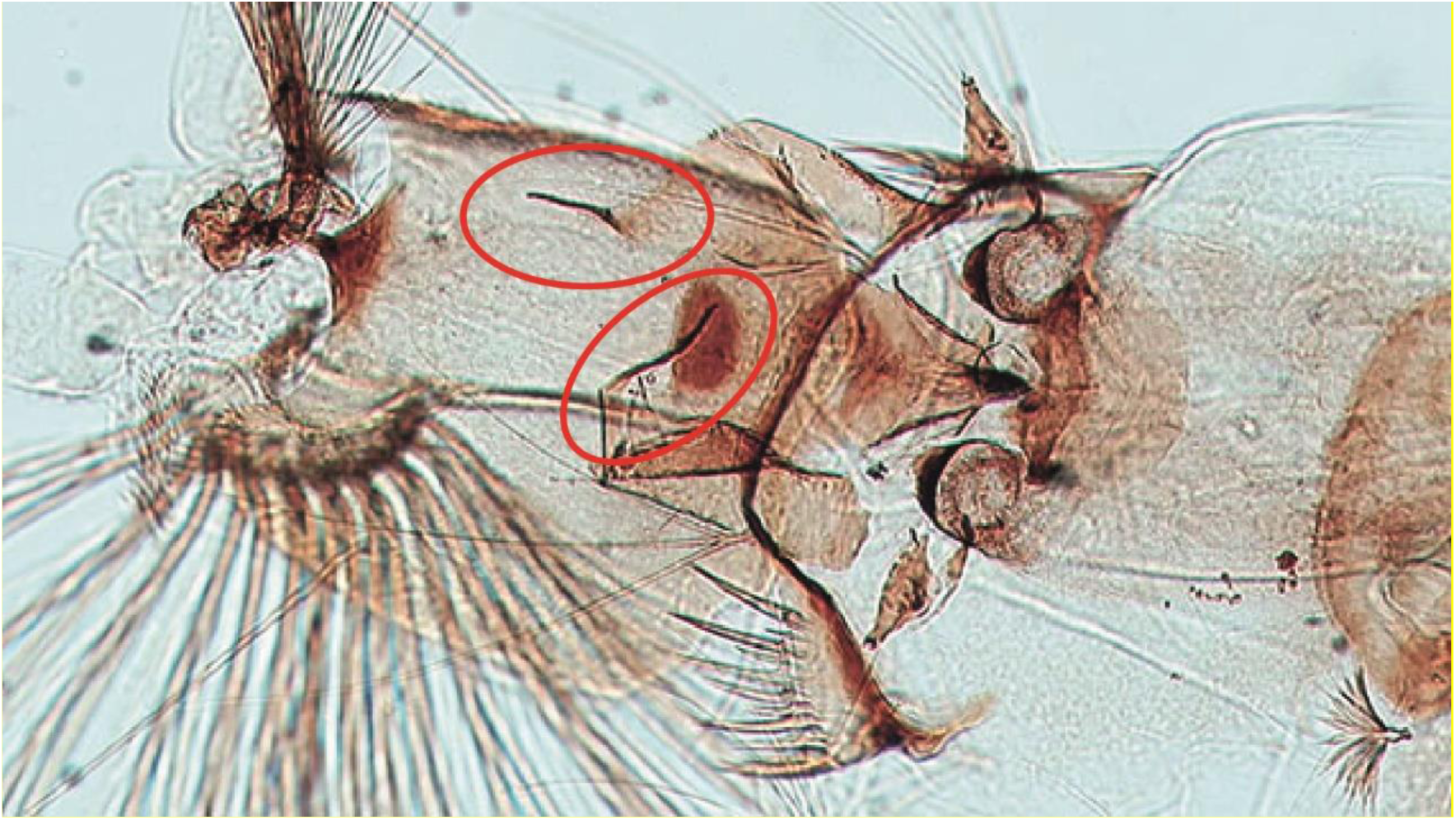
Last segments of an *An. pseudopunctipennis* larva collected in Lincoln Park, Cochabamba (collection code: LTA-2024-01-140:001), showing the spiracular apparatus. The two ellipses highlight the characteristic "black tails," a unique feature of *An. pseudopunctipennis* larvae.

### Specimen preservation and storage

All collected samples from the project and mounted specimens, including those with *An. pseudopunctipennis,* have been deposited in the collections of the Medical Entomology Laboratory (LEMUMSS) at the Universidad Mayor de San Simón in Cochabamba, Bolivia.

### Maxent habitat suitability model for *An. pseudopunctipennis*

A habitat suitability analysis for *An. pseudopunctipennis* was performed using an ecological niche modeling approach with the Maxent algorithm (version 3.4.4) ^15, 16^, adhering to established best practice guidelines ^17–19^. The analysis comprised the following sequential steps:

#### Occurrence data

Occurrence data for *An. pseudopunctipennis* in Bolivia were sourced from the VECTOBOL database which aggregates georeferenced collection points throughout the country, with vector species identified by scientists. A dataset of 218 points, where *An. pseudopunctipennis* larvae were collected, was extracted from the database.

#### Environmental predictors

The initial set of the 19 bioclimatic variables (BIO1 to BIO19) from the WorldClim ver. 2.1 database ^20^ at a resolution of 30 arcsecond represented the various candidate predictors that are potentially relevant for the geographic distribution of *An. pseudopunctipennis* (data available at: https://www.worldclim.org/data/worldclim21.html). This dataset is representative of annual and seasonal means variations of temperatures and precipitations averaged over a 30-year period (1970-2000).

#### Selection of relevant environmental predictors

When constructing models with collinear variables, there is an elevated risk of overfitting and overparametrization ^21^. The reduction of relevant variables was therefore achieved through statistical analysis, with a focus on retaining those variables that held biological significance for *An. pseudopuntipennis*. The WorldClim variables were submitted to a VIF (Variance Inflation Factor) analysis using the R-package *usdm* ^22^. In the package, the two functions *vifcor* and *vifstep* were used with recommended thresholds of 0.7 ^23^ and 5 ^24, 25^ respectively. The function *vifcor*, first find a pair of variables which has the maximum linear correlation (greater than the threshold), and exclude one of them which has greater VIF. The procedure is repeated until no variable with a high correlation coefficient greater than the threshold with other variables remains. The function *vifstep* calculate VIF for all variables, exclude one with highest VIF (greater than threshold), repeat the procedure until no variables with VIF greater than the threshold remains. The two functions were applied to the pixel values of all 19 bioclimatic raster layers within the area defined by a convex hull of 150 km around each occurrence records and an altitude <3 300 m. This convex hull delineates the training area for the Maxent model. The two analyses consistently selected the four variables BIO2 (mean diurnal range*; i.e.* mean monthly (Temperature max Temperature min)), BIO3 (isothermality, *i.e*., BIO2/BIO7*100, where BIO7 is the temperature annual range, *i.e.,* the temperature maximum of the warmest month – the temperature minimum of the coldest month), BIO5 (the maximum temperature of the warmest month), and BIO18 (the precipitation of the warmest quarter). Correlations among these four variables were all below 0.7 (Table 1), and the VIF values were 1.85, 2.24, 2.75 and 1.44, respectively.

**Table 1.**
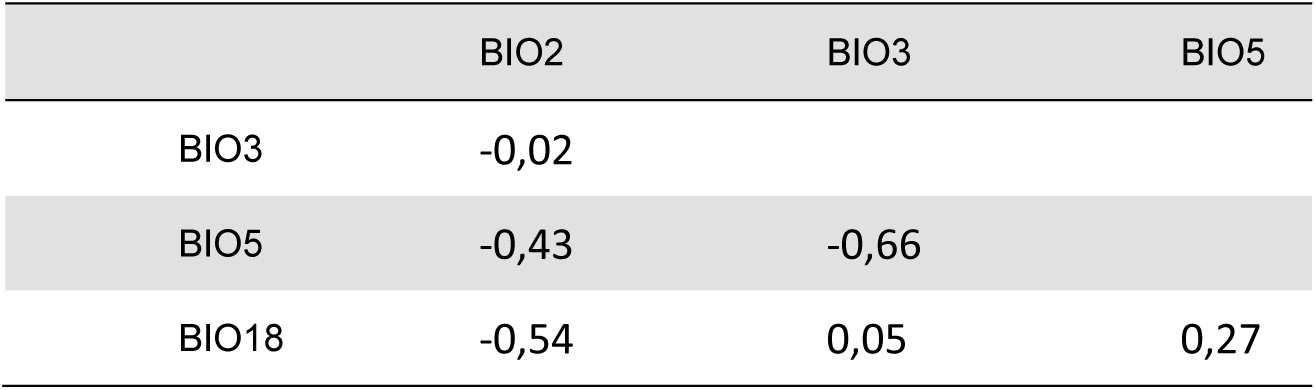
Correlation matrix between variables BIO2, BIO3, BIO5 and BIO18.

#### Occurrence data cleaning

In a clean occurrence dataset, data are not spatially autocorrelated. Indeed, spatial autocorrelation within the occurrence dataset indicates a lack of independence among geographically proximate observations, and can potentially affect the model outcomes. Spatial autocorrelation among the 213 *An. pseudopunctipennis* occurrences was evaluated through the analysis of a multivariogram using the *variogmultiv* function from the R-package *adespatial*, and considering the four selected bioclimatic variables. The analysis revealed significant autocorrelation at 2 arc-minute distance (approximately 4 km) (Figure 2A). Consequently, the initial occurrence dataset of 213 points was thinned using the Rpackage *spThin* to remove geographically close points, resulting in a final dataset of 77 collection points.

**Figure 2.**
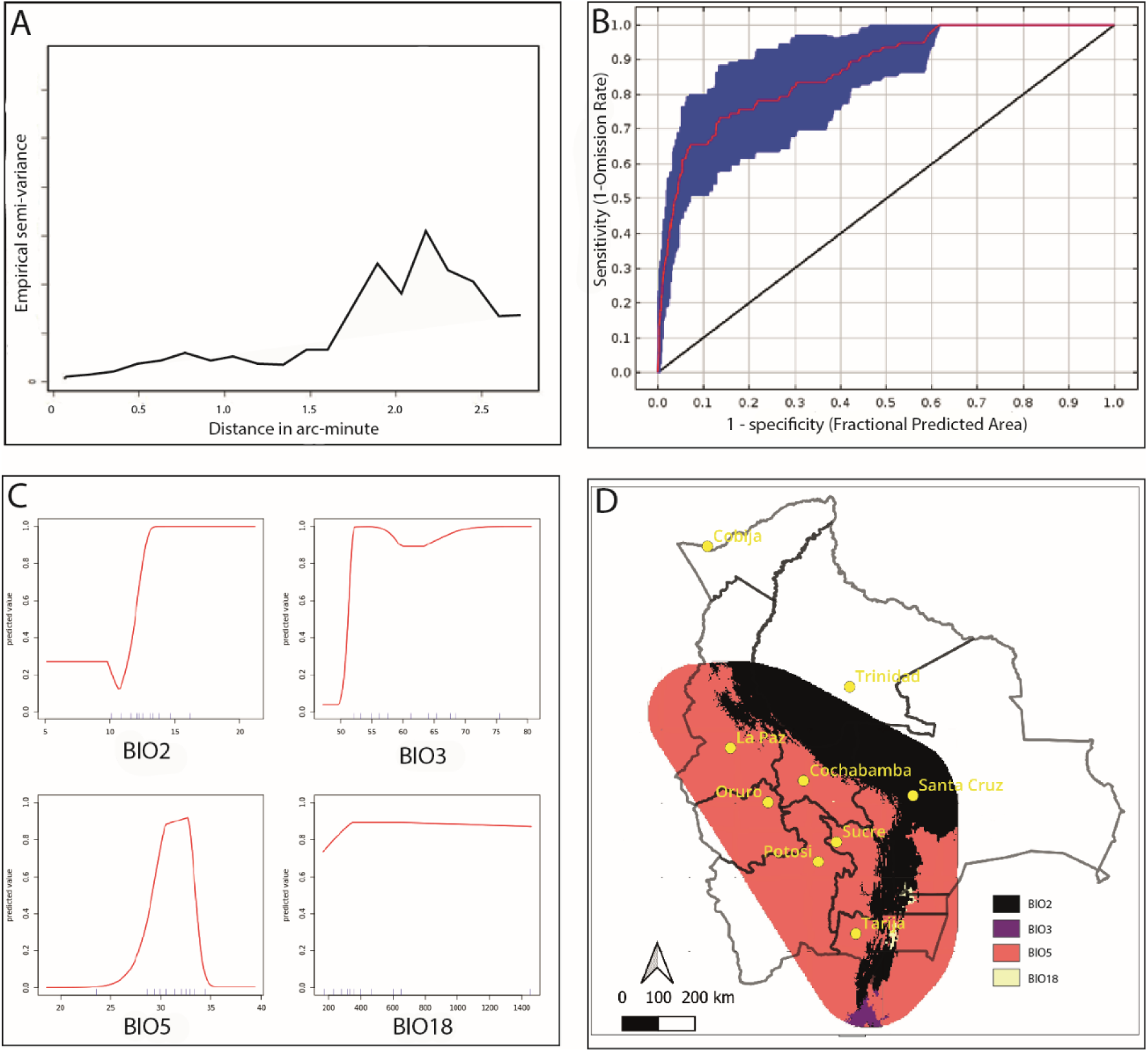
Maxent modeling. (A) Multrivariogram based on 213 occurences of An. *pseudopunctipennis* and the bioclimatic WorldClim variables BIO2, BIO3, BIO5 and BIO18. (B) Receiver Operating Characteristic (ROC) curve from Maxent output. (C) Response of An. *pseudopunctipennis* to environmental variables BIO2, BIO3, BIO5 and BIO18. (D) Map of limiting factors in the training area of the Maxent model.

#### Maxent Model building

The Maxent model was created using the R-package *sdm* ^26^ by randomly selecting 80% of the 77 occurrences for the training data set, and the four selected climatic variables. Model performance was evaluated based on significance (partial *ROC*, with 500 iterations and 20% of data for bootstrapping) and omission rates (*OR*, with parameter *E*=5%). The final model was constructed as an ensemble, averaging the 8 top-performing models selected based on their AUC (Area under Curve) and TSS (True Skill Statistics) values from 100 replicates, employing recommended bootstrapping techniques ^27^. A jackknife process was executed to evaluate the relative importance of single explanatory variables included in the model. The number of occurrence data for *An. pseudopunctipennis* (n=77) is sufficient to construct a robust Maxent model, enabling accurate predictions of habitat suitability across Bolivia ^28^.

## Results

### *An. pseudopunctipennis* collection site and voucher specimens

The entomological survey of the UMSS-DB.07 project was conducted during the rainy season in March and April 2024 and the dry season in August, September, and October 2024. A total of 251 larval samples were collected (162 during the rainy season and 89 during the dry season), of which 32 were considered potentially suitable for *An. pseudopunctipennis* larvae, characterized by clear water from springs, ponds with algae, and river margins with algae. However, only one was site was found positive for *An. pseudopunctipennis* and was located in Lincoln Park, in the center of Cochabamba city. The larval habitat (Lat: -17,37020; Long: -66,16911; Alt: 2 470 m), sampled on April 16, 2024, consisted of a pond with an inner stone perimeter and a cement bottom, covering an area of 184 square meters with a depth of one meter. The water was clear, with very slow flow from a small spring, and abundant green algae of the genus *Rhizoclonium* concentrated in the central part of the pond (Figure 3 A, B and C). Excess water drained into a small stream. The site exhibited characteristic features of suitable habitats for this species, including optimal water physico-chemical quality, low current, and the presence of green algae. *Culex brethesi* shared this breeding site with *An. pseudopunctipennis*.

**Figure 3.**
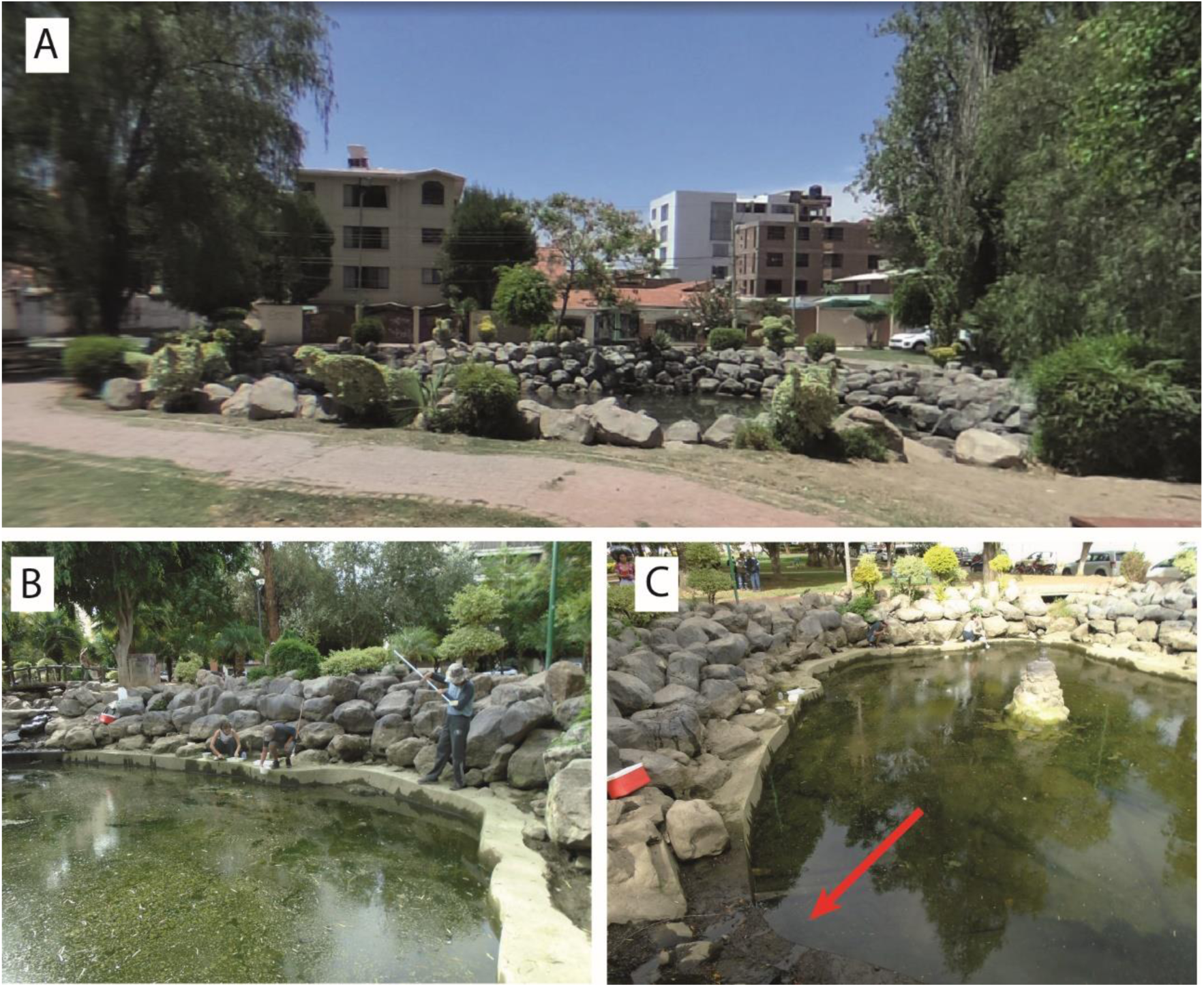
Larval habitat of An. *pseudopunctipennis* in Lincoln Park, Cochabamba city. (A) General view, (B) Larval habitat with *Rhizoclonium* algae, (C) Larval habitat showing clear water from a small spring (red arrow).

The field sample containing *An. pseudopunctipennis* is registered under No. LTA-2024-01-140:001 in the VECTOBOL database. Voucher specimens of *An. pseudopunctipennis* have been deposited in the specimen collection of the entomology laboratory (LEMUMSS) of the Universidad Mayor de San Simón, Cochabamba, with identification codes matching the ID prefix of the collected samples to ensure traceability within the laboratory’s collection system. The assigned voucher codes are LTA-2024-01-140:001 to LTA-2024-01-140:004.

### Maxent model of *An. pseudopunctipennis* potential distribution

Figure 4 shows the resulting habitat suitability map from Maxent modeling, focusing on Cochabamba city. The inset map in Figure 4 displays the model’s projection at the country level. The corresponding raster file, along with other Maxent outputs, is available at Zenodo (https://doi.org/10.5281/zenodo.17826929).

**Figure 4.**
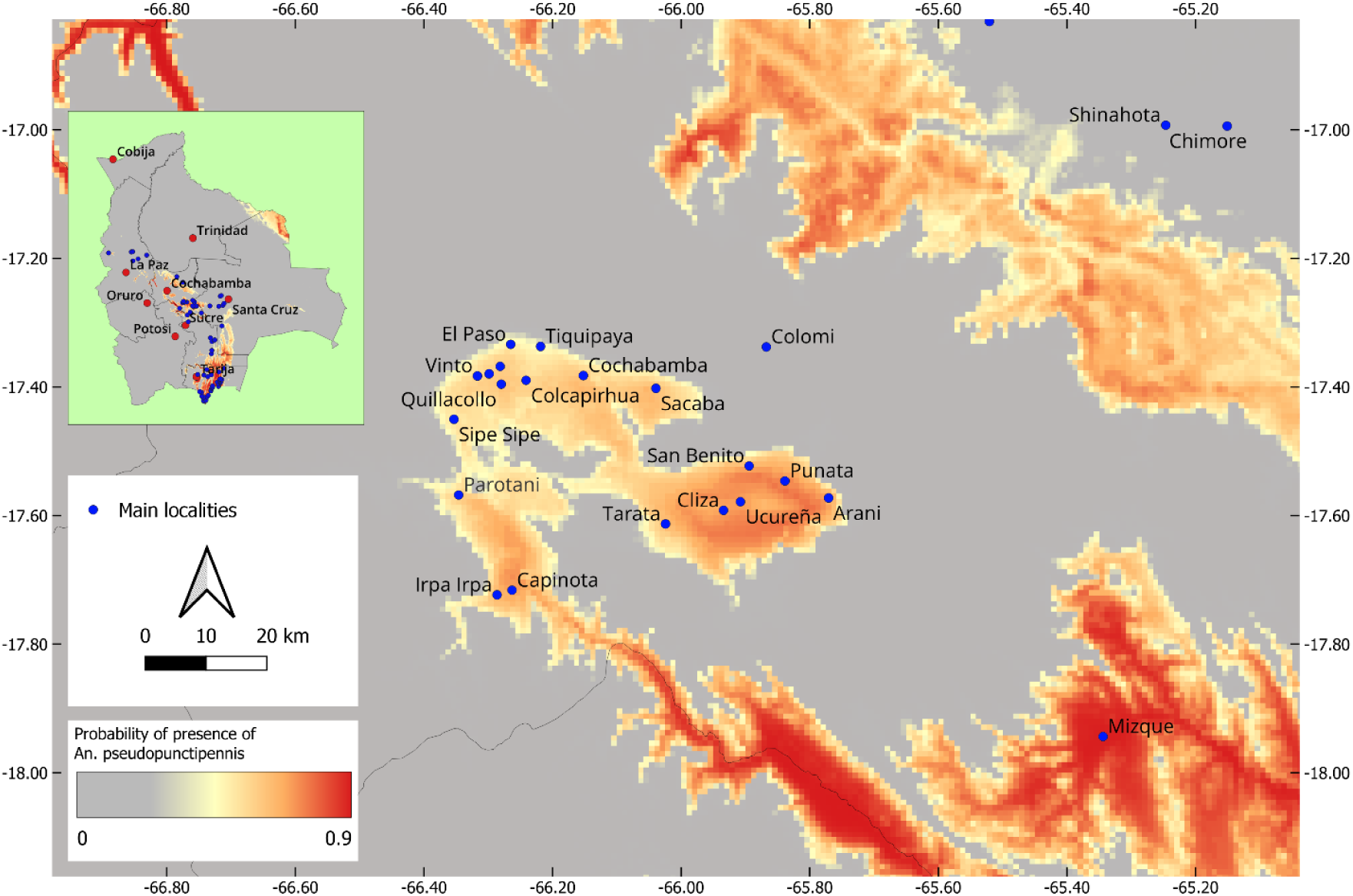
Maxent modelling: map of the probability of presence of An. pseudopunctipennis in the geographical area of Cochabamba city. The inset map shows the model projection at the country level with the An. pseudopunctipennis collecting points (blue dots) and the department capitals (red dots).

Table 2 presents the percentage contribution and permutation importance of the environmental variables BIO2, BIO3, BIO5 and BIO18. The percentage contribution measures the extent to which each variable contributed to the final model, with higher values indicating a greater influence on the prediction of habitat suitability. Permutation importance assesses the model’s dependency on each variable; higher values indicate that shuffling the variable’s values leads to a significant decrease in model performance, suggesting that the variable is critical to the model’s predictive ability. Values highlight the predominant influence of variable BIO5 (maximum temperature of the warmest month) in the Maxent model predictions, followed by BIO2, with BIO3 also playing a noticeable role, albeit to a lesser extent, while BIO18 has minimal influence.

**Table 2.**
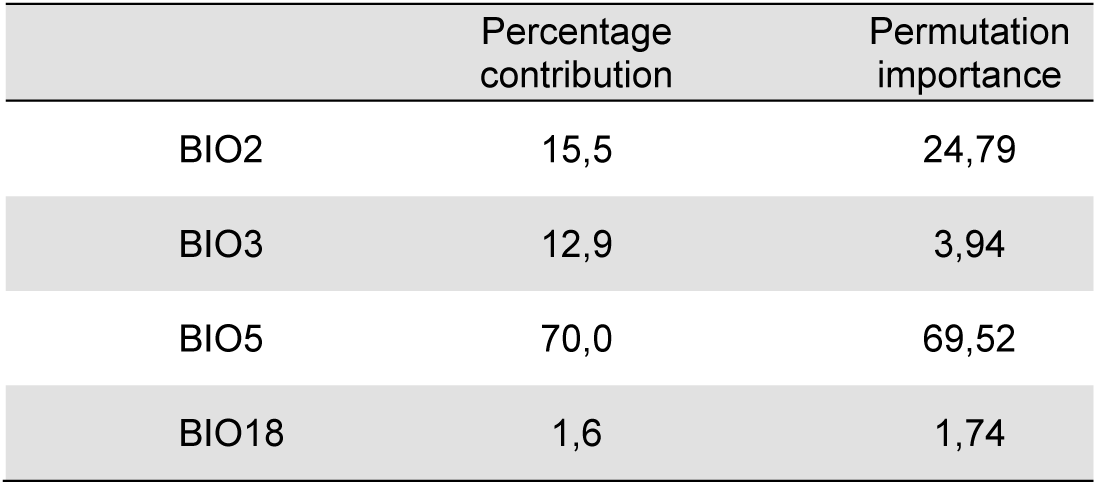
Percentage contribution and permutation importance of environmental variables BIO2, BIO3, BIO5 and BIO18.

**Table 3.**
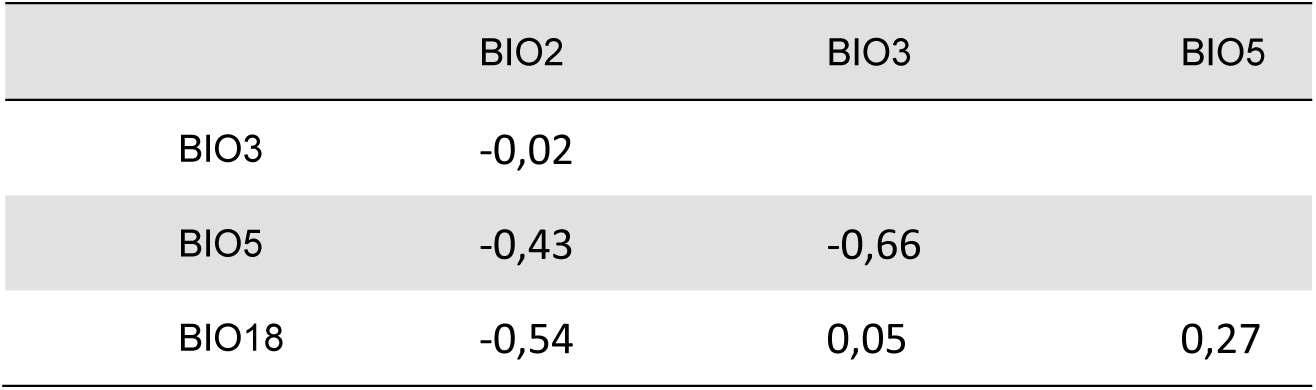
Correlation matrix between variables BIO2, BIO3, BIO5 and BIO18.

**Table 4.**
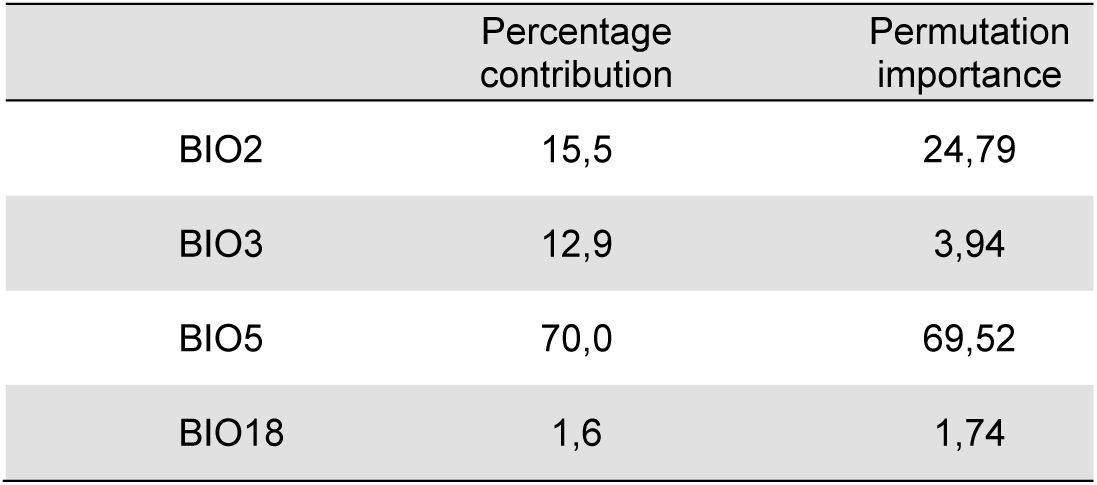
Percentage contribution and permutation importance of environmental variables BIO2, BIO3, BIO5 and BIO18.

The Receiver Operating Characteristic (ROC) curve is given in Figure 2B, with the AUC value of 0.87, significantly greater than the null model (AUC=0.68), with a TSS value >0.7, demonstrating the effectiveness of the Maxent model. The variables’ response curves indicate how habitat suitability changes with respect to individual environmental variable used in the Maxent model and are given in Figure 2C. In the model training area, the bioclimatic variable BIO5 is the limiting factor in the western region of Bolivia, including the Cochabamba area. In contrast, BIO2 is the limiting factor in the east, while BIO3 and BIO18 act as limiting factors in specific patches in the southeast (Figure 2D).

As indicated by the model, the bioclimatic variables, particularly BIO5, delineate a suitable habitat for *An. pseudopunctipennis* in the Cochabamba Valley.

## Discussion

Larvae of *An. pseudopunctipennis* are found in diverse habitats, but they frequently inhabit sun-exposed freshwater stream pools containing filamentous green algae such as *Spirogyra*, *Oedogonium*, *Cladophora*, *Closterium*, and *Enteromorpha*. These environments, characterized by clear, shallow, and stagnant water, provide oviposition substrates for gravid adult females as well as food and shelter for the larvae ^29–31^. In Cochabamba city, the larval habitat where *An. pseudopunctipennis* was collected matches this description, featuring clear, quasi-stagnant water and abundant green algae of the *Rhizoclonium* genus.

*Anopheles pseudopunctipennis* is recognized as a malaria vector within its geographical range and has long been suspected to be a vector in Bolivia. Concrete evidence confirming its role as a vector in Bolivia was recently documented in 2008 ^32^, based on wild-caught individuals from Andean valleys. However, its discovery in Cochabamba is not cause for alarm. For an epidemic to occur, the vector must reach a certain density relative to the human population. The current survey of mosquito larval habitats in Cochabamba city and neighboring areas indicate very few suitable breeding sites, which are often contaminated, hindering establishment of the species. Th species was found in only one site, in very low density. *Anopheles pseudopunctipennis* has not disappeared from the area but is now limited in abundance. It should be regarded more as an ecological curiosity, possibly nearing extinction due to increasing urbanization destroying its natural breeding sites and water contamination. However, the potential geographic distribution of *An. pseudopunctipennis*, as predicted by the Maxent model, suggests that Cochabamba city and neighboring areas offer suitable habitat for this species. Furthermore, the altitude of the area falls within the species’ distribution range where it can be found up to 3 200 m ^33, 34^. The VECTOBOL initiative has recorded data points up to 2 800 m in localities such as Tuntunani and Mollebamba in the La Paz Department, and 2 700 m in Aiquile in the Cochabamba Department. Therefore, if sufficient larval habitats become available in the area, such as shallow riverbanks, residual river pools, shallow irrigation canals with slow streams, and resurgences, with clear water and filamentous algae ^35, 36^, the species could proliferate during favorable seasons, as was likely the case in the city of Santa Cruz de la Sierra in 1997 ^37^.

The origin of the collected larvae in Cochabamba city is unknown. The most parsimonious hypothesis is that there is a very small population of *An. pseudopunctipennis* that can sustain itself there, with individuals potentially migrating from neighboring areas (within a few kilometers) when climatic conditions are suitable. These habitats, however, have yet to be identified in subsequent surveys focusing on canals and river ponds in the close area. The Maxent model also identified a more remote and adequate area delineated by the localities of San Benito, Punata, Arani, Tarata, and Cliza, southeast of Cochabamba (Figure 4). As this area is more rural and likely less affected by pollution, the presence of suitable and populated larval habitats is plausible. However, according to SEDES Cochabamba, only a few malaria cases were reported in 2023 in the Depatrment, with no distinction between imported and autochthonous cases. While the possibility of rare autochthonous malaria cases cannot be entirely ruled out, the low abundance of *An. pseudopunctipennis* suggests that it is unlikely to pose a significant risk for malaria transmission in the Cochabamba city area.

## Conclusion

The discovery of *An. pseudopunctipennis* in the city of Cochabamba serves as a reminder that almost 90 years ago, the area was a malarial zone where epidemics occurred. Currently, the disease is absent from the area, with only a few rare annual cases, often imported from other malarial regions. Thus, the presence of this species in Cochabamba is more of a scientific curiosity than a significant public health threat. Given the urbanization of the area and the consequent destruction of the species’ natural habitats, it is unlikely that *An. pseudopunctipennis* will re-establish itself in the city. However, it would be interesting to survey close areas that remain relatively rural, such as the formerly malarial areas of Parotani and Capinota to the southwest of Cochabamba, as well as the Cliza-Punata-Tarata region to the southeast.

## Acknowledgments

The authors thank the students who participated in the UMSS-DB.07 project, assisting with field sampling and laboratory processing of the samples: Sofia Y. Quispe Sanchez, Cristian Zubieta Sempertegui, Samuel Caero Heredia, Marcelo Quevedo Ledezma, Ariana Bustamante Michel, Alex Flores Herrera, and Maria F. Rodriguez Echalar.

## Statement on AI usage

The authors declare that no artificial intelligence (AI) tools were used to generate, edit, or write any part of this article. All content was produced solely by the authors.

